# *Plasmodium falciparum* Replication factor C subunit 1 is involved in genotoxic stress response

**DOI:** 10.1101/2020.05.26.118224

**Authors:** O Sheriff, Y Aniweh, Soak-Kuan Lai, HL Loo, S. K Sze, PR Preiser

## Abstract

About half the world’s population is at risk of malaria, with *Plasmodium falciparum* malaria being responsible for the most malaria related deaths globally. Antimalarial drugs such as chloroquine and artemisinin are directed towards the proliferating intra-erythrocytic stages of the parasite, which is responsible for all the clinical symptoms of the disease. These antimalarial drugs have been reported to function via multiple pathways, one of which induces DNA damage via the generation of free radicals and reactive oxygen species. An urgent need to understand the mechanistic details of drug response and resistance is highlighted by the decreasing clinical efficacy of the front line drug, Artemisinin.

The replication factor C subunit 1 protein is an important component of the DNA replication machinery and DNA damage response mechanism. Here we show the translocation of PfRFC1 from an intranuclear localization to the nuclear periphery indicating an orchestrated progression of distinct patterns of replication in the developing parasites. PfRFC1 responds to genotoxic stress via elevated protein levels in soluble and chromatin bound fractions.

Reduction of PfRFC1 protein levels upon treatment with antimalarials suggests an interplay of replication and DNA repair pathways leading to cell death. Additionally, mislocalization of the endogenously tagged protein confirmed its essential role in parasites’ replication and DNA repair. This study provides key insights into DNA replication, DNA damage response and cell death in *plasmodium falciparum*.

**Importance:** Frontline drugs have been found to induce DNA damage in the human malaria parasite *Plasmodium falciparum.* The genotoxic stress response in *Plasmodium* and the interplay between DNA damage repair, replication and activation of programmed cell death pathways remains largely undescribed. This study shows a distinct pattern of localization of PfRFC1 during replication and DNA repair. PfRFC1 responds to genotoxic stress with an increase in protein expression. Interfering with the RFC complex formation or mislocalization of PfRFC1 is associated with disrupted genotoxic stress response. Additionally, a reduction of PfRFC1 protein levels is observed upon treatment with antimalarial drugs or under apoptosis like conditions, highlighting the role of DEVD/G like motif in mediating programmed cell death in these parasites. This study sheds light on the role of PfRFC1 in differentially responding to replication, genotoxic stress and programmed cell death in *Plasmodium* parasites.

## Introduction

*Plasmodium falciparum*, the main causative agents of malaria and responsible for most malaria related human deaths, has a digenetic lifecycle involving distinct developmental forms. The intraerythrocytic parasite has a haploid genome which is subjected to exogenous and endogenous insults arising via normal cellular processes such as; replication errors, heme degradation and exposure to antimalarial drugs like artemisinin, mefloquine and chloroquine^1–3^. Artemisinin resistant strains have also been associated with down regulation of DNA replication genes during ring^4^, trophozoite and schizont stages^5^.

The replication factor C (RFC) complex has been identified to be differentially regulated in artemisinin resistant parasites^6^. The Replication Factor C is a heteropentameric complex of five subunits (RFC1/2/3/4/5) conserved both in structure and function^7–10^. The RFC functions as a clamp loader of the proliferating cell nuclear antigen (PCNA) sliding clamps and as a RFC-PCNA-DNA complex bringing about DNA synthesis^11^. The RFC complex subunits contain conserved motifs termed RFC boxes I-VIII. These are responsible for DNA binding, PCNA interactions, DNA replication and RFC complex maintenance^12^. The RFC box I, present in the N-terminal of the larger RFC1 subunit, shows high homology to prokaryotic DNA ligases and BRCT domains^13^ and is involved in DNA binding^14^ The four small RFC subunits from human and yeast align with the central part of the larger RFC1 subunit denoted by boxes II-VIII^10^ and belong to the AAA+ (ATPases Associated with diverse cellular Activities) superfamily of ATPases. The boxes II-VII are also important for DNA binding and loading of PCNA^15,16^. The interaction of RFC1 with PCNA has also been shown to be essential for the synthesis of repair templates during the two types of DNA repair mechanisms; DNA excision repair and double strand break repair (DSBR)^17^. Among the RFC subunits, the regions of the amino acid similarity exist in the N-terminal half of the protein, while the C-terminal regions of the subunits are unique and are required for the formation of the RFC complex^18,19^. RFC1 is also involved in apoptosis via a conserved putative caspase-3 cleavage site at the C-terminus of the DNA binding region. Fragments of RFC1 released upon protease activity inhibit DNA synthesis and promote apoptosis^20,21^.

The *Plasmodium* parasites activate both excision repair and DSBR mechanisms to remove DNA insults via the replacement of the DNA damage induced histone modifications^22^. PfRFC1 involved in nucleotide excision repair is reported to be up-regulated by MMS induced genotoxic stress in *Plasmodium falciparum* parasites^22^. The ability of a parasite-cell-free lysate to repair apurinic/apyrimidinic sites revealed that *Plasmodium* parasites perform repair via the long patch base excision repair (BER) pathway^23^.This is unlike the mammalian and yeast systems which resort to a short one-nucleotide based repair. Other members of this pathway such as; PfFEN1, PfPCNA1 and PfPCNA2 have been reported to be expressed at higher levels in response to DNA damaging agents in *Plasmodium* parasites^24^. Homologous recombination repair; a form of DSBR, has been demonstrated to be the most effective DNA repair mechanism employed by the parasite to overcome deleterious double strand breaks in the DNA. Bioinformatics as well as homology modelling tools have been used to show the conservation of most of the components of the nucleotide excision repair mechanism^25,26^. Furthermore, experimental evidence suggests the presence of a functional alternate non-homologous end joining DSBR mechanism^27,28^. Transcriptome studies have also highlighted the activation of numerous DNA repair mechanisms upon treatment with genotoxic agents^22^. However, there remains a lack of understanding of the interplay between these molecules as well as the functional characterization of the individual components.

Despite its importance in yeast and human cellular processes, the large subunit of the RFC complex in *Plasmodium* intra-erythrocytic cell cycle remains uncharacterized. In this study, PfRFC1 in *Plasmodium falciparum* was endogenously tagged and its localization was identified to be dynamic in asexual lifecycle of the parasite. Further, immunoprecipitation assay confirmed its interaction with the sliding clamp loader PCNA1 and identified other members of the complex. Treatment of *Plasmodium falciparum* parasites with genotoxic agents showed elevated protein levels of PfRFC1 and change in its localization. An N-terminal truncation of PfRFC1containing the RFTS, when expressed in addition to the native PfRFC1 affected recovery from genotoxic stress, highlighting the essential role of RFC1 in DNA damage repair in *P. falciparum.* Finally, conditional mislocalization of RFC1 confirmed its essential role in cell cycle progression of the asexual stages of the parasites as well as recovery from genotoxic stress.

## Materials and Methods

### Parasite culture and synchronization

*Plasmodium falciparum* strains were cultured in human red blood cells donated within 30 days prior to usage. Parasites were cultures in RPMI1640 media supplemented with 50mg/l gentamicin, 2g/l sodium bicarbonate, 0.25% Albumax II, and 0.1 mM hypoxanthine. The cultures were maintained in 1-2% parasitemia at 37°C under microaerophilic conditions. Cultures were synchronized using 5% D-sorbitol or by 68% percoll purification of schizonts followed by 5% D-sorbitol treatment post invasion. Synchronization of the parasites were verified by morphology of the parasites via giemsa stained thin blood smears.

### Plasmids construction, Parasite transfections and integration conformation

To generate the SLI (selection-linked integration) knock-in constructs a 551bp C-terminal fragment excluding the stop codon of PfRFC1 was PCR amplified from the *P. falciparum* genomic DNA. This fragment was inserted between the NotI/AvrII in the pSLI-2×FKBP-GFP^29^ for pSLI-PfRFC1-GFP, and NotI/MluI sites after replacing the GFP tag with a codon optimized 3xHA tag in the pSLI-2×FKBP-GFP plasmid for pSLI-PfRFC1-HA constructs using primers F.RFC1_NOT1, R.RFC1_HA_MLU1 and R.RFC1_GFP_AVR2 (Table S1).

The construct for the pDC2-RFC1Δ2-HA cell line was generated by firstly PCR amplifying a 939bp N-terminal fragment between the Not1/Mlu1 site of pSLI-RFC1-HA to get the construct pSLI-RFC1Δ2_HA using the primers F.RFC1Δ2_HA_NOT1 and R.RFCIΔ2_HA_MLU1 (Table S 1). This construct was used as a template to amplify the truncated RFC1 gene with the 3xHA tag using the primer F.RFC1Δ2_HA and R.RFC1Δ2_HA (Table S1).This fragment was inserted between the AvrII/XhoI site of the modified pDC2 plasmid^30^.

pSLI-PfRFC1-HA and pSLI-PfRFC1-GFP plasmids were used to generate PfRFC1-HA and PfRFC1-GFP cells respectively by the following method. Synchronized ring stage parasites were transfected with 100 – 200 μg plasmids purified by NucleoBond Xtra Midi EF midiprep kit (Macherey-Nagel) by electroporation using Bio-Rad laboratories gene pulser X cell^31^. 2.5nM WR99210 (Jacobus Pharmaceuticals, Princeton, NJ) was added 6-8hours post transfection for the pSLI construct and with blasticidinS (Invivogen) at 2 μg/ml for the pDC2 plasmid. The media was changed daily for 5 days subsequently and drug selection was maintained until parasites were observed in the giemsa smears. Tagged protein expression was verified by western blotting prior to conducting assays.

For the pSLI plasmids, selection linked integration was performed as described^32^ on parasites obtained from the first round of selection with WR99210 carrying the episomal plasmid. These were subjected to 400 μg/ml G418 (Gibco). Parasites obtained were tested for correct integration by the following PCR primers for the endogenous tagging of PfRFC1 (Table S1). F_RFC1 and N_MLU1_PSLI confirmed the junction upstream of the site of integration, while primers FJ8C_PARL and R_RFC 1 _UTR confirmed the region downstream of the plasmid at the site of integration. F_RFC1 and R_RFC1_UTR confirmed the absence of the unmodified locus in the selected parasites (Table S1). Confirmed PfRFC1-GFP cells were subsequently transfected with pLyn-FRB-mCherry-nmd3-BSD plasmid expressing the plasma membrane mislocalizer under blasticidinS selection^32^. mCherry and GFP expressing parasites were enriched by counting 2 million cells on a BD FACSAria. Appropriate gating of cells was established using 3D7 parental parasites. These parasites were cultured under standard culture conditions prior to assays.

### Co-Immunoprecipitation

Immunoprecipitations were performed on late stage trophozoite parasites by percoll enrichment. Enriched parasites were first treated with 0.5mM DSP (dithiobis[succinimidyl propionate], Thermo Scientific Pierce) for 30 min. The reaction was quenched with excess of 25mM Tris-HCl pH7.5 in PBS for 10 min. The parasites were subsequently released from the RBC via 0.05% saponin (Sigma) in incomplete RPMI (RPMI1640 medium without AlbumaxII). The pellet fraction was then solubilized in 10 volumes of RIPA buffer (Thermo Fisher Scientific) with benzonase nuclease (EMD Millipore) and Halt protease inhibitor cocktail (Thermo Fisher Scientific) for 30min on ice. The clarified lysates were precleared with Protein A conjugated magnetic beads (Pierce) for 1 hour with rotation at 4C to remove non-specific protein binding. The precleared fraction was split equally between Anti-HA Magnetic Beads (Pierce) and Anti-c-Myc Magnetic Beads (Pierce) to immunoprecipitate HA tagged proteins and nonspecific proteins respectively. These were incubated at 4C with rotation overnight. The IP beads were washed at least 5 times in cold RIPA and proteins were eluted by heating in non-denaturing loading buffer according to manufacturer’ s recommendation and then subjecting the eluted fraction to denaturation using Dithiothreitol (DTT) and heat.

### Genotoxic agent treatment of *Plasmodium* parasites

HU (Hydroxyurea) (Sigma) and MMS (Sigma) were used for DNA-damage studies by treating synchronized late stage trophozoite (~36 hour) *P.falciparum* at 5% parasitemia. parasites were treated with MMS (0.005%) or HU (10mM) for 6 hour at 37°C in normal culture conditions in line with earlier studies on related replication proteins^24^. The parasites were retrieved for immunofluorescence assay. The parasites were released from erythrocytes with 0.015% saponin in PBS, followed by multiple washes. Saponin extracted parasites were subjected to subcellular fractionation as described below or lysate preparation followed by western blot analysis. LiCor compatible secondary antibodies such as IRDye 800CW and IRDye 680LT goat anti-rabbit, goat anti-mouse and goat anti-rat were used in according to manufacturer’ s recommendation. Western blots were processed using the standard protocol by LiCor and imaged using a LiCor odyssey CLx imager and software. The data was analyzed and reported as mean±SD (n=3). Statistical analysis was performed using Microsoft excel and the student’s t test was used to measure differences between means and p≤0.05 was marked significant.

### Drug treatment of Plasmodium parasites

Chloroquine was dissolved in water and filter sterilized to obtain working solutions of 1mM which were made fresh and stored at 4’C in the dark. CQ treatment was performed at 1x IC_50_ (30nM), 10x IC_50_ (300nM), 100xIC_50_ (3μM) concentrations. Artesunate (ART), an artemisinin derivative was dissolved in dimethylsulfoxide (DMSO). ART treatment of the parasites was performed at concentrations of 1xIC_50_ (2nM) and 10xIC_50_ (20nM). Parasitized erythrocytes were incubated for 6h prior to harvesting for western blot analysis. Staurosporine (ST, Sigma-Aldrich) stock solution (1 mM) was prepared by dissolving the drug in filtered DMSO and stored at −20°C. 100 mM of working ST solution was prepared before each experiment by diluting the stock solution with RPMI. Concentrations of Staurosporine (ST) at 1, 2, 5 μM were used. Infected erythrocytes were treated for time and concentrations required prior to being washed twice with culture medium for assays. Controls for necrosis were generated by incubating parasites with 1-0.1%(W/v) of sodium azide for the required duration. Vehicle controls with DMSO or water were also utilized.

### In vitro parasite survival assay

To determine the survival of parasites after treatment with genotoxic agents, synchronized late stage trophozoites (~36 hour) at 1% parasitemia were subjected to concentrations of MMS ranging from 0.00005% to 0.005% for 6h. After the 6 hour incubation period the parasites were washed multiple times in RPMI medium and returned to fresh medium. Parasite recovery was measured ~18 hours after MMS washout. The parasites were stained with the nuclear stain Hoechst 33342 (Sigma) and parasitemia was measured via the Attune NxT Flow Cytometer (Thermo Fisher Scientific) or LSRFortessa™ X-20 (BD Biosciences).

### IFA (Immunofluorescence Assay), antibodies & microscopy

Synchronized *Plasmodium falciparum* infected erythrocytes were smeared onto glass slides and air dried. The slides were subsequently fixed for 5 min with cold methanol at −20°C. These slides were rehydrated in 1x PBS at room temperature for 15 min followed by blocking for 1 hour in 1x PBS containing 3% BSA (Bovine serum albumin, Sigma). The slides were then incubated for 1 hour at room temperature with 1xPBS+3% BSA with respective antibodies. Primary antibodies used in this study were anti-EXP2 Rabbit^33^, anti-HA Rat (Roche), anti-Histone H3 Rabbit (Abcam), anti-GFP mouse (Abcam) and anti-H3K9Me3 Rabbit (Abcam). The slides were washed for 15 min with 1x PBS/0.1% Tween20 post primary antibody treatment followed by 1 hour incubation at room temperature with 1x PBS+3%BSA containing secondary antibodies anti rabbit IgG-Alexa 488 or anti rat IgG-Alexa 594 (all from Jackson ImmunoResearch). The nuclear stain DAPI (2μg/ml) was then applied followed by washes with 1xPBS/0.1% Tween20 at room temperature. The coverslips were mounted using Fluoromount-G (southernbiotech). Slides were visualized on a Nikon Eclipse Ti fluorescent microscope with a Nikon Plan Apochromat Lambda 100X Oil objective. Pictures were taken using an Andor Zyla sCMOS camera and analysed using ImageJ 1.52n. For quantitative imaging, the images were captured with a Zeiss LSM710 confocal microscope equipped with an Airyscan detector (Carl Zeiss) using a Plan-Apochromat 100x/1.46 oil objective. These images were processed using imageJ 1.52n and Adobe Photoshop CS6.

### Total RNA extraction and Real time PCR analysis

*Plasmodium falciparum* strains; W2mef, Dd2, Gb4, 3D7, K1 and NF54 well cultivated and D-Sorbitol synchronized. The parasites were harvested in 8-hour time difference across the 48 hour life cycle. The harvested parasites were stored in TRIzol (Ambion/Life Technologies) pending RNA and DNA extraction. The parasites stages were homogenized in TRIzol with DNA and total RNA was extracted using the Direct-Zol RNA MiniPrep Plus kit (Zymo Research) according to manufacturer’s protocol. Expression of mRNA transcripts for PfRFC1 gene analysis were carried out using the Luna Universal One-Step RT-qPCR Kit (New England Biolabs, Inc.), on a QuantStudio 5 Real-Time PCR System (Applied Biosystems), using manufacturer’ s recommendations. The primers sets used in the assay are set1 and/or set2 for purposes of confirming the expression level (Table S1). The seryl-tRNA synthetase gene expression was used as the endogenous control. The Ct values generated from the expression analysis were converted to expression levels using the 2^-ΔΔCt^ formula. Data were analyzed with Microsoft Excel and GraphPad software (v.7).

### Subcellular Fractionation of Parasites

The detergent soluble and the insoluble fraction were prepared by adopting an existing protocol^34^. Parasites extracted with 0.015% saponin were lysed in Buffer A [20 mM HEPES (pH7.9), 10 mMKCl, I mM EDTA, 1 mM EGTA, 0.3% NP-40 and 1mM DTT] and Halt protease inhibitor cocktail (Thermo Fisher Scientific) with incubation on ice for 5 min. The insoluble nuclear fraction was pelleted down at 2700 x g and the soluble fraction was recovered. The pellet fraction was washed multiple times with buffer A followed by lysate preparation and western blot analysis. The efficiency of fractionation was confirmed by antibodies Anti-Histone H3 (Millipore) (marker for insoluble nuclear fraction) and Anti-PfAldolase Rabbit (GenScript) (marker for soluble fraction).

### Protein complex characterization by immunoprecipitation tandem mass spectrometry

Immunoprecipitated and eluted proteins from each fractions were separated on 12% SDS-PAGE at 50 V and protein bands were visualized by staining with imperial protein stain (Pierce). The gel lanes corresponding to each of the fractions (HA beads and Control beads) were cut into 2 separate slices, then de-stained, and the proteins reduced by using dithiothreitol (DTT) and alkylated by iodoacetamide (IAA). The proteins were cleaved by overnight digestion in porcine trypsin (Sequencing Grade Modified, Promega, Wisconsin). The tryptic peptides were extracted by using 5% acetic acid in 50% acetonitrile and vacuum-dried by speedvac. The vacuum concentrated peptides were reconstituted in 0.1% formic acid (FA) and 3% ACN for LC-MS/MS analysis in the Q-Exactive Hybrid Quadrupole-Orbitrap mass spectrometer, coupled with the UltiMateTM 3000 RSLCnano System (Thermo Scientific Inc, USA). The peptides were first concentrated with a Nano-Trap Columns 75-100 μm I.D. x 2 cm (Thermo Scientific, USA) and then separated on a Dionex EASY-Spray 75 μm x 10 cm column packed with PepMap C18, 3 μm, 100 Å (Thermo Fisher Scientific, USA). The mobile phase buffers used were 0.1% formic acid (A) and 0.1% formic acid in ACN (B) and a 60 min gradient was used for peptide separation. The samples were ionized and injected into the Q-Exactive mass spectrometer with an EASY nanospray source (Thermo Fisher Scientific, Inc.) at an electrospray potential of 1.5 kV. A full MS scan (350–1,600 m/z range) was acquired at a resolution of 70,000, with a maximum ion accumulation time of 100 ms. Dynamic exclusion was set as 30 s. The HCD spectral resolution was set to 35,000. Automatic gain control (AGC) settings of the full MS scan and the MS2 scan were 3E6 and 2E5 respectively. The top 10 most intense ions above the 5,000 count threshold were selected for fragmentation in higher-energy collisional dissociation (HCD), with a maximum ion accumulation time of 120 ms. Isolation width of 2 was used for MS2. Single and unassigned charged ions were excluded from MS/MS. For HCD, the normalized collision energy was set to 28% and the under fill ratio was defined as 0.3%.

### Database searching

The raw data generated for each sample were analyzed using the Proteome Discoverer (PD) 1.4 software (Thermo Scientific, San Jose, CA). Protein identification was done by mapping against a customized protein sequence database combined from UniProt *Homo sapiens* proteome, the *P. falciparum* 3D7 proteome in PlasmoDB 13.0 and common contaminant database (http://maxquant.org/contaminants.zip and ftp://ftp.thegpm.org/fasta/cRAP/crap.fasta), using the SequestHT and Mascot search engines. The Proteome Discoverer’s workflow included an automatic target-decoy search tactic along with the Percolator to score peptide spectral matches from both Mascot and SequestHT searches to estimate the false discovery rate (FDR). The Percolator parameters are set to maximum delta Cn = 0.05; target FDR (strict) = 0.01; target FDR (relaxed) = 0.05, validation based on q-value. The search parameters also included full trypsin digestion with a maximum of two missed cleavage and precursor mass tolerance and fragment mass tolerance was set at 10 ppm and 0.02 Da respectively. Carbamidomethylation (+57.02) at cysteine was set as fixed modification, oxidation (+15.99) at methionine, deamidation (+0.98) at asparagine and glutamine. Precursor ion area was used for protein quantitation.

Protein identifications were considered valid if ≥2 unique peptide sequences were detected. Statistically significant peptide matches corresponding to specific protein hits in bait IP reactions (or alternatively, peptide enrichment was compared with the control) were collated into tables ordered based on peptide enrichment (Table S2). Proteins known to be typical contaminants were excluded from the analysis. For charting, the number of peptides displayed represents the total number of peptide matches across all replicates and experiments.

## Results

### RFC1 localizes dynamically through intraerythrocytic developmental cycle

In order to confirm the identity and conservation of *Plasmodium falciparum* RFC1 (PfRFC1, PF3d7_0219600), its amino acid sequence was compared with human (HsRFC1) and yeast (ScRFC1) to reveal a sequence identity of ~28% (Fig. 1A). PfRFC1 also shared the important functional conserved boxes I-VIII regions (Fig. 1B). The N terminus of human and yeast RFC1 possess a 20 amino acid replication factory targeting sequence (RFTS) consisting of a PCNA interacting protein motif (PIP)^35^ and a stretch of positively charged residues. This RFTS found at the N-terminal region is highly conserved across the species and was recessed by ~90 amino acids within the N-terminal of PfRFC1 (Fig. 1C).

**Fig. 1:**
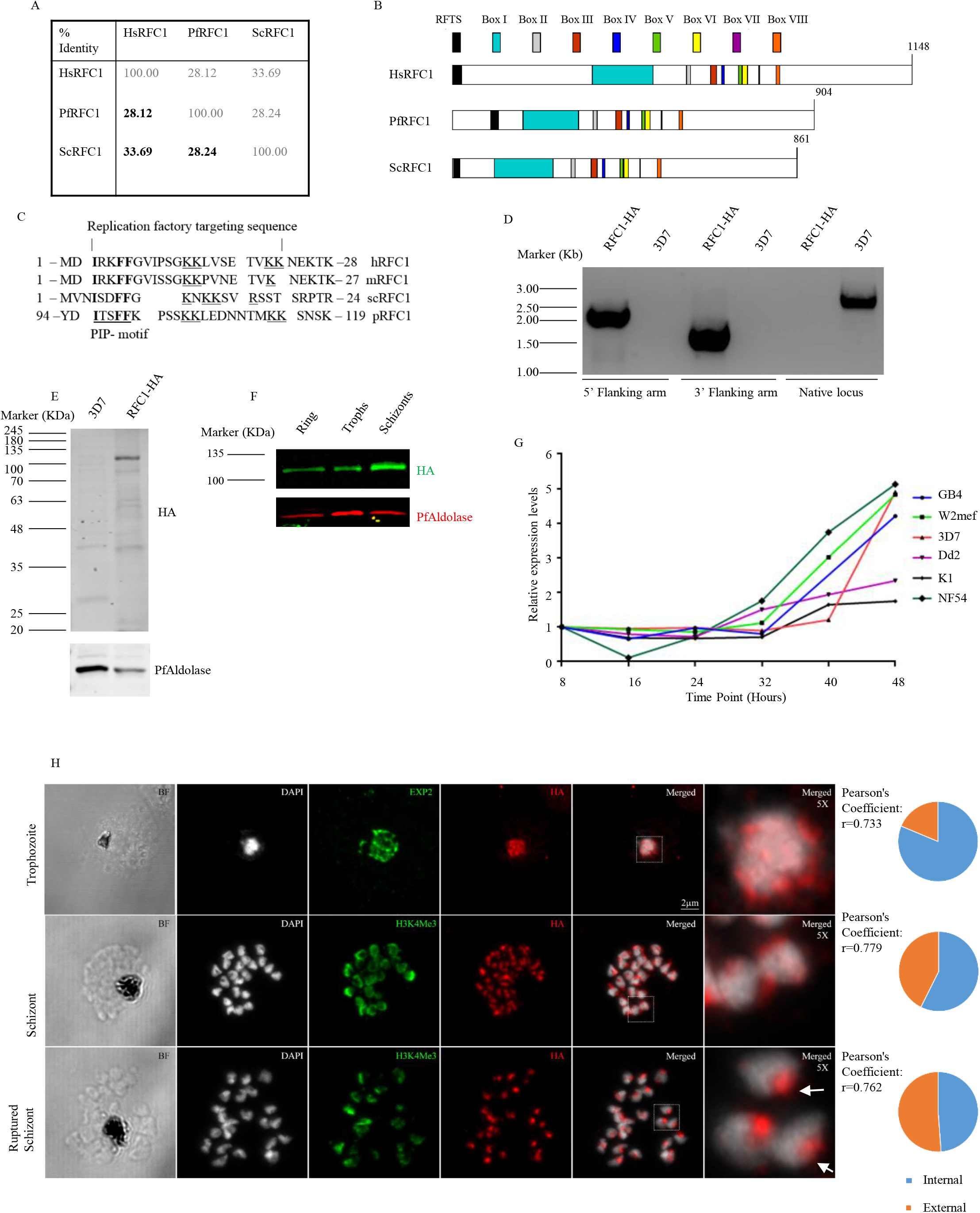
Endogenously tagged PfRFC1 localizes dynamically throughout intraerythrocytic developmental cycle. **(A)** PfRFC1 shows low sequence identity to homologs in human (HsRFC1) and yeast (ScRFC1). **(B)** PfRFC1 contains all the conserved boxes in their corresponding order with **(C)** an N-terminal extension prior to the replication factory targeting sequence (RFTS) in comparison to the RFTS in the RFC1 of Human(hRFC1), mouse (mRFC1) or yeast (ScRFC1). **(D)** PfRFC1 was endogenously tagged with a 3xHa tag and integration was verified via PCR. **(E)** Tagged PfRFC1 was identified by anti-Ha antibody in the Parasite fraction. **(F)** PfRFC1 (green) was identified in the ring, trophozoite and schizont stages and PfAldolase was used as a loading control (Red). **(G)** Real-Time PCR show variability in the stage specific expression levels of the RFC1 transcript across the various laboratory strains. **(H)** Super resolution microscopy on immunofluorescence samples of trophozoite and schizont, and ruptured schizont stages show specific nuclear localization (Pearson’s Coefficient: r=0.733 for trophozoites; r=0.779 for schizonts; 0.762 for ruptured schizonts measured for DAPI and HA co-localization). The segmented schizonts shows perinuclear localization while the ruptured schizont show a single punctae in the nucleus (white arrow). DAPI (White), PfEXP2/ H3K4Me3 (Green), and HA (Red). 5x Magnified insets of representative trophozoite, schizont and ruptured schizonts nuclei are provided. The pie-charts reflect the PfRFC1 signal % within and outside the nucleus respectively of at least 3 individual cells.

The protein expression profile and localization was studied by generating transgenic parasites expressing PfRFC1 tagged with a triple haemagglutinin tag (PfRFC1-3xHA) in its endogenous loci via SLI (selection-linked integration)^29^. The successful integration was confirmed by PCR (Fig. 1D) and the protein was detected by western blotting in the parasite fraction (Fig. 1E). PfRFC1 was more abundant in the schizont stages (Fig. 1F). RT-PCR performed on RNA extracted every 8h over the 48h life cycle of 6 laboratory strains of *P.falciparum* revealed a basal level of expression in the early stages of the cell cycle with an increase of 2-5 fold observed in the schizont stages (Fig. 1G). seryl-tRNA synthetase gene expression was used as the endogenous control. Interestingly, the two multidrug resistant strains K1 and Dd2 showed only ~2 fold stage dependent up-regulation in schizont stages (48 hours). The increased transcription of PfRFC1 observed in most strains correlated with increased DNA synthesis in the parasite leading up to schizogony and is in line with previous transcription studies^36,37^ and reports showing increased replisome protein levels at the trophozoite and schizont stages^38^.

As RFC1 has been shown to be recruited to the replication foci in other organisms, immunofluorescence microscopy was used to establish the localization of PfRFC1 in PfRFC1-3xHA parasites. Consistent with the western blot data, immunofluorescence shows the presence of PfRFC1 in the ring, trophozoite and schizont stages. Co-localization with the DAPI stained nucleus indicated nuclear localization of PfRFC1 (Fig. S1). Labelling the parasite periphery with anti-Exp2 showed that PfRFC1 is found parasite internal and is located in numerous punctate nuclear foci within the trophozoite nucleus (white) marked by DAPI (Fig. 1H). These foci are consistent with the observation of replication factories at sites of active replication in these stages^24,39^. In segmented schizonts, PfRFC1 while still co-localizing with DAPI, is predominately observed at the periphery of the segmented nuclei (Fig. 1H). Further, egressed merozoites show PfRFC1 in a single punctae at the edge of each nuclei (Fig. 1H, white arrows). Quantification of these nuclei across the stages was performed for multiple cells to determine the presence of the signals of PfRFC1 within and outside the nuclei marked by DAPI. The Pie-charts show that the signal of PfRFC1 is predominantly inside the nucleus in the trophozoite stages while progressively migrating outside the nucleus as the parasite matures to schizonts and free merozoites where they are observed in the nuclear periphery. The localization of PfRFC1 is significantly altered between the trophozoite and schizont stages where the total PfRFC1 signals within the nucleus are 81.3±8.07 % in trophozoites and 56.63±0.79 % in schizont (p<0.05). The Pearson’s Coefficient indicated in the figure (r=0.733 for trophozoites; r=0.779 for schizonts; 0.762 for ruptured schizonts) measures the colocalization of HA with DAPI, and reflects the co-localization of the PfRFC1 signal with the nuclear stain respectively (Fig. 1H). The pattern of localization of PfRFC1 was compared with that of PfH3K4Me3 one of the most abundant histone marks in the genome of *P.falciparum* and postulated to be associated with intergenic regions. PfH3K4Me3 has been reported to be localized in a horseshoe shaped pattern at the nuclear periphery of the parasites representing sites of active translation^40,41^. The signals of H3K4Me3 localize with the sites of PfRFC1 in schizont (pearson’s coefficient of 0.711) and diverge in merozoites stages (pearson’s coefficient of 0.459) correlating with the end of replication (Fig. 1H).

### RFC1 reveals a complex associating with PCNA1

In order to identify the interacting partners of PfRFC1, Co-immunoprecipitation (CoIP) was performed on PfRFC1-3xHA late stage trophozoites and the eluted proteins were identified by mass spectrometry. Successful Co-IP was confirmed by gel staining (data not shown) and by western blot to detect the enrichment of PfRFC1 in the HA beads (Fig. 2A). PfPCNA1 was also found to be enriched in the HA beads confirming the interaction of PfRFC1 at the replication foci. PfAldolase, a cytoplasmic protein and PfHistone3, a nuclear marker, were used as controls and were not identified on the bead fractions by western blotting. Mass spectrometry of the IP fraction identified a total of 59 *P.falciparum* proteins with at least 2 unique peptides detected in HA bead sample and 6 of these were also detected in the control bead sample of the trophozoite stage IP (Table S2). Among these proteins heat shock proteins and annotated ribosomal proteins were excluded as common contaminants leaving behind 39 proteins. Importantly, the most highly abundant interacting partners with ≥ 4 peptides enriched on the HA beads (Fig. 1B) represent the expected subunits of the Replication factor c consisting of proteins PfRFC1-5. Nuclear proteins such as PfRAN, DNA replication licensing factor PfMCM2, and topoisomerase I were also enriched in the IP. PfPCNA1 although detected by Western blot was only identified by one unique peptide by mass spectrometry. Numerous exported, glycolytic, proteolysis and RNA binding proteins such as PfAlba1were also identified in the IP and are likely non-specifically or weakly associated with the PfRFC1 interactome. Metabolic pathways enrichment analysis of the Co-immunoprecipitated proteins showed an abundance of proteins involved in oxidative stress and DNA replication and repair (Fig. 1C) highlighting the role of PfRFC1 in these processes.

**Fig. 2:**
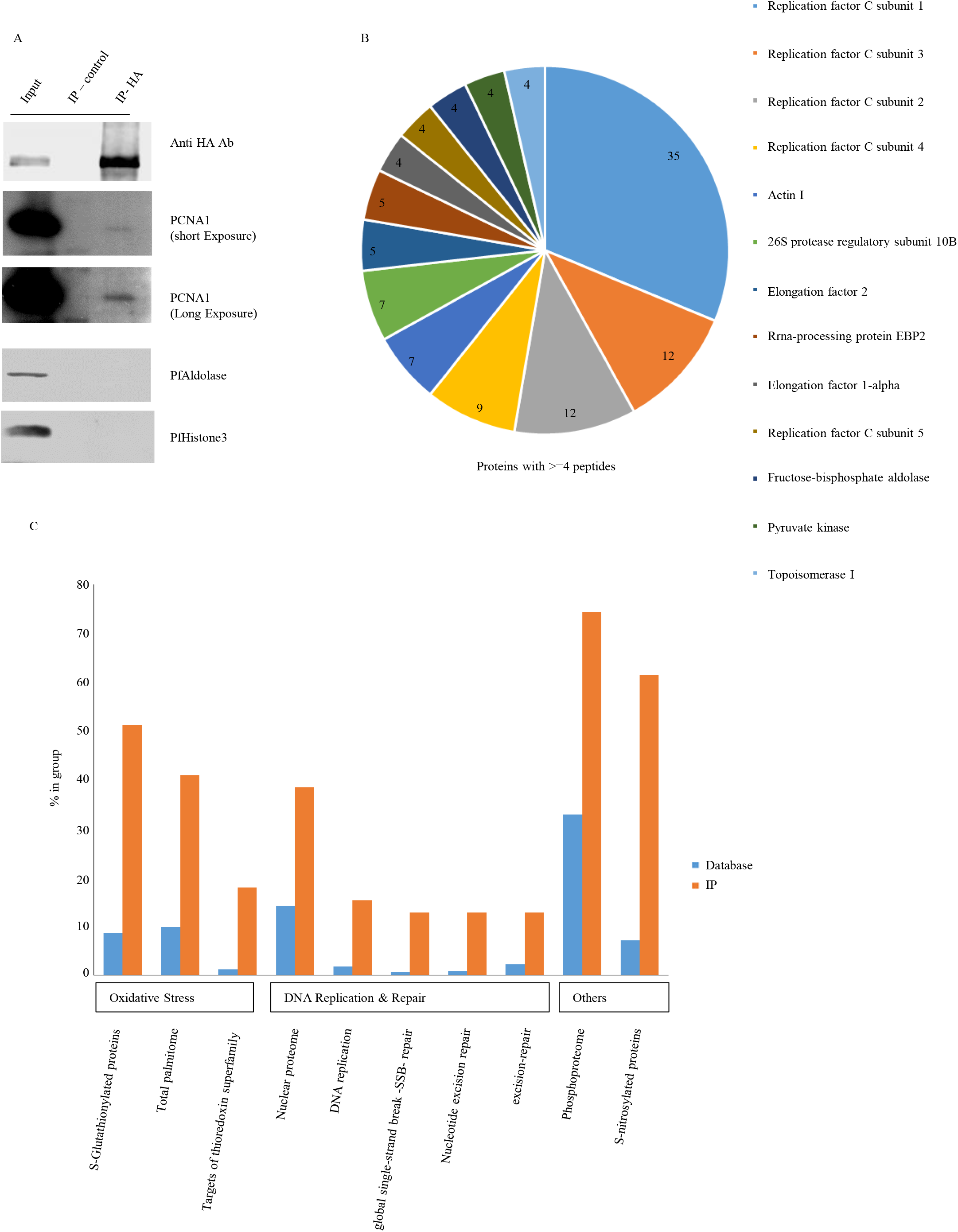
IP of PfRFC1 confirms a complex associating with PfPCNA1. **(A)** Immunoprecipitation of PfRFC1 was performed and the various samples were loaded on a SDS-Page and subjected to western blotting as follows. The lane one contains the input parasite lysate, lane 2 contains the eluted fraction from control beads and the lane 3 contains the eluted fractions from the HA beads. The blots were probed with anti HA antibody to confirm the specific immunoprecipitation of PfRFC1 at the expected size, PfAldolase and PfHistone 3 were used as a loading controls. PfPCNA1 was identified in the HA bead fraction which also contains the enriched PfRFC1. PfPCNA1 was observed to be specifically present in the HA beads although in low abundance as seen via the short and long exposure of the blot.**(B)** The Pie-chart represents proteins with ≥4 peptides detected to be enriched on the HA beads as compared with the control beads. These proteins are enriched in the components of the Replication factor C. Number of peptides enriched are mentioned within the pie chart for each protein **(C)** A malaria metabolic pathway enrichment profile of the immunoprecipitated proteins shows enrichment of components involved in oxidative stress, DNA replication and repair as compared with the database at plasmodb.org (Release 44).

### PfRFC1 is stimulated upon DNA damage

We investigated the involvement of PfRFC1 in DNA repair as the IP of PfRFC1 indicated the presence of PfPCNA1 enriched in the PfRFC1 IP sample. To evaluate the impact of genotoxic stress on PfRFC1, synchronized early trophozoite stages were treated with MMS (0.005%) or HU (10mM) for 6h in line with earlier studies on related replication proteins^24^. In *P.falciparum,* MMS has been verified via comet assays to cause DNA damage leading to elevated transcripts of repair components PfRAD51,PfRAD54, PfRFC1^22,42^ and elevated protein levels of PfRAD51, PfPCNA1 and PfPCNA2^24^. MMS, an alkylating agent is responsible for stalling of DNA synthesis in the S-phase via single and double strand breaks^43^, while HU targets ribonucleotide reductase and induces genomic instability by arresting replication fork progression due to the depletion of dNTPs^44,45^. Western blot analysis of equal fractions of treated parasites showed an increased protein expression of PfRFC1 by 3.4± 0.57 fold upon MMS treatment (n=3, p=0.014) and 2.85 ± 0.51 upon HU treatment (n=3, p=0.023) (Fig. 3A, B) as compared to control treatment. PfAldolase, a constitutively expressed protein served as a control. In parallel, parasites treated with MMS and HU were fractionated into soluble and chromatin bound fractions. These samples were analyzed by western blotting and probed with anti HA antibody to detect PfRFC1, PfAldolase for the soluble fraction and PfHistone3 for the chromatin bound fraction. Three independent experiments confirmed the significant increase of PfRFC1 levels upon treatment with genotoxic agents (Fig. S2 A, B). The increase of RFC1 in the whole cell lysates upon DNA damage was also reflected in its enrichment in both the soluble and the chromatin bound fraction highlighting its role in the response to genotoxic agents.

**Fig. 3:**
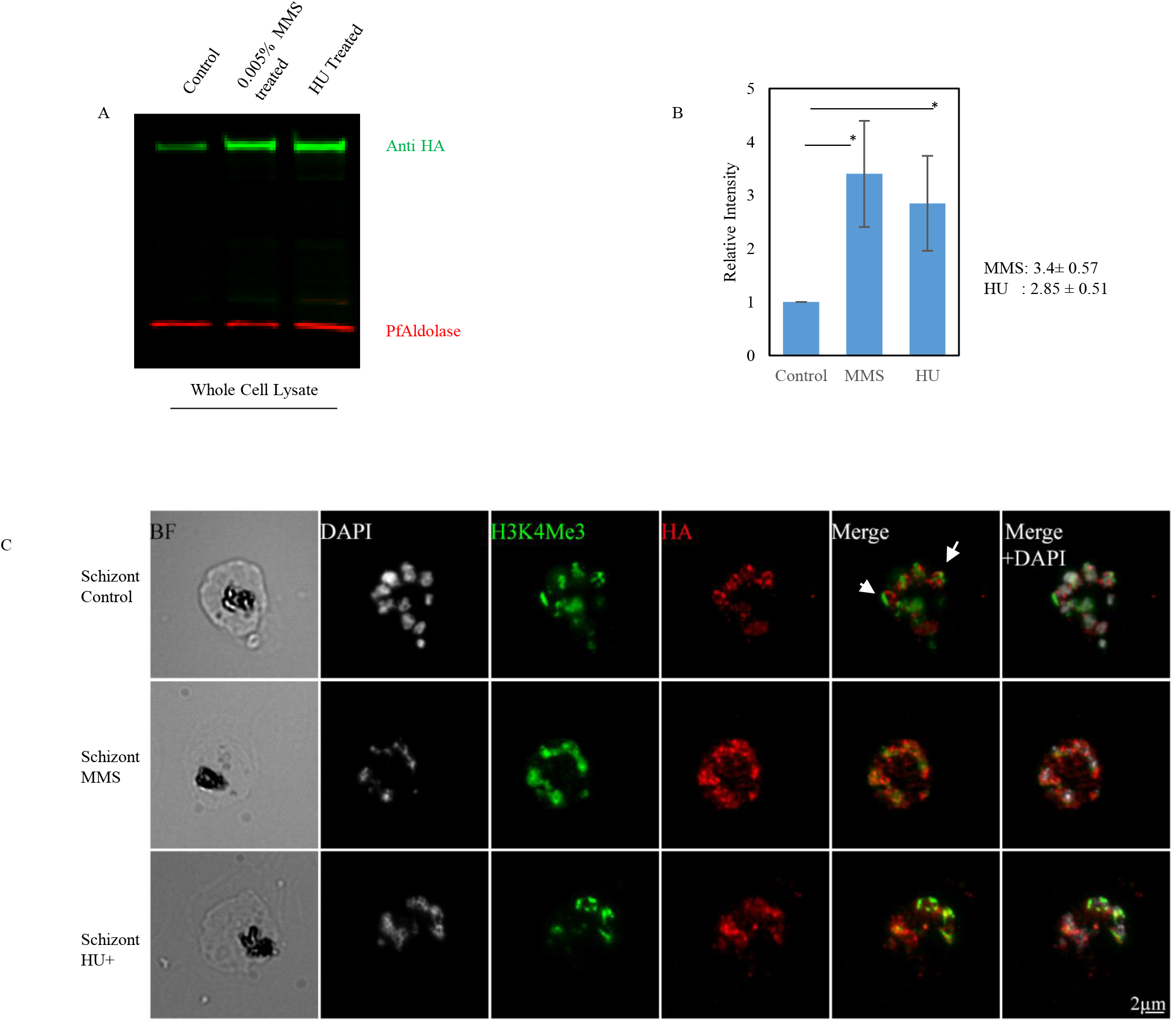
PfRFC1 is stimulated upon DNA damage. **(A)** Endogenously tagged PfRFC1 expressing trophozoite cells were treated with control (Lane 1), MMS (Lane 2) or Hydroxyurea (Lane 3) and western blotting of the whole parasite lysates was performed. The levels of PfAldolase were used as the loading control to determine PfRFC1 levels using anti HA antibodies. **(B)** Three independent experiments were performed and the densitometric analysis of RFC1 was normalised to that of PfAldolase and represented graphically. The results show means ±S.E.M (n=3,p=0.014 for MMS and p=0.023 for HU).**(C)** Immunofluorescence was performed on the developing schizonts from the above treatments and probed with anti HA (Red), Anti H3K4Me3 (Green) and stained the nucleus with DAPI (White). The Perinuclear staining in the control treated samples for PfRFC1 was not observed in the cells subjected to genotoxic stress. White arrows indicate regions where PfRFC1 and H3K4Me3 do not co-localize.

Immunofluorescence assay on fixed PfRFC1-3xHA schizont parasites after MMS or HU treatment showed that while in the untreated control PfRFC1 was located at the nuclear periphery, this changed to a more dispersed location within the nucleus in the parasites subjected to genotoxic stress (Fig. 3C). PfRFC1 in trophozoite stage parasites was observed via immunofluorescence to form punctate patterns throughout the nucleus in control as well as treated samples as expected of its role in replication and repair. (Fig. S2C). The signal of PfRFC1 shows partial co-localization with PfH3K4Me3 as observed previously (Fig. 3C). It however did not completely localize with PfH3K4Me3 highlighting the compartmentalization of PfRFC1 from these regions in late stage schizonts (Fig. 3C, white arrows).

### Critical role of PfRFC1 in DNA damage recovery

To assess the role of PfRFC1 upon genotoxic stress in greater detail we co-expressed a truncated fragment of PfRFC1. The N-terminal of PfRFC1 including the RFTS and BRCT domain was tagged with a 3xHA tag and overexpressed in *P.falciparum*^29^. The selected parasites replicated at a rate comparable to the controls indicating no significant effect on replication. The truncated PfRFC1 (PfRFC1 Δ2-HA) lacks the AAA+ATPase domain as well as the RFC1 C-terminal homology domain (Fig. 4A). The exclusion of the C-terminal RFC1 homology domain prevents the assembly of the RFC1-5 complex while continuing to interact with PCNA1 via the RFTS impairing excision repair. The expression of PfRFC1Δ2-HA was verified with a band at 41.1 KDa via western blotting on the parasite lysates (Fig. 4B). Since PfRFC1Δ2-HA contains the BRCT domain, reported to be involved in DNA binding, the ability of the truncated protein to be associated with the chromatin fraction was investigated (Fig. 4C).The truncated PfRFC1 Δ2-HA was observed in both the soluble and the chromatin bound fraction confirming its ability to associate with chromatin. Immunofluorescence assay on the parasites expressing the truncated PfRFC1Δ2-HA also showed nuclear localization (Fig. 4D) while the signals remained diffuse unlike the punctate pattern observed for the full length PfRFC1.

**Fig. 4:**
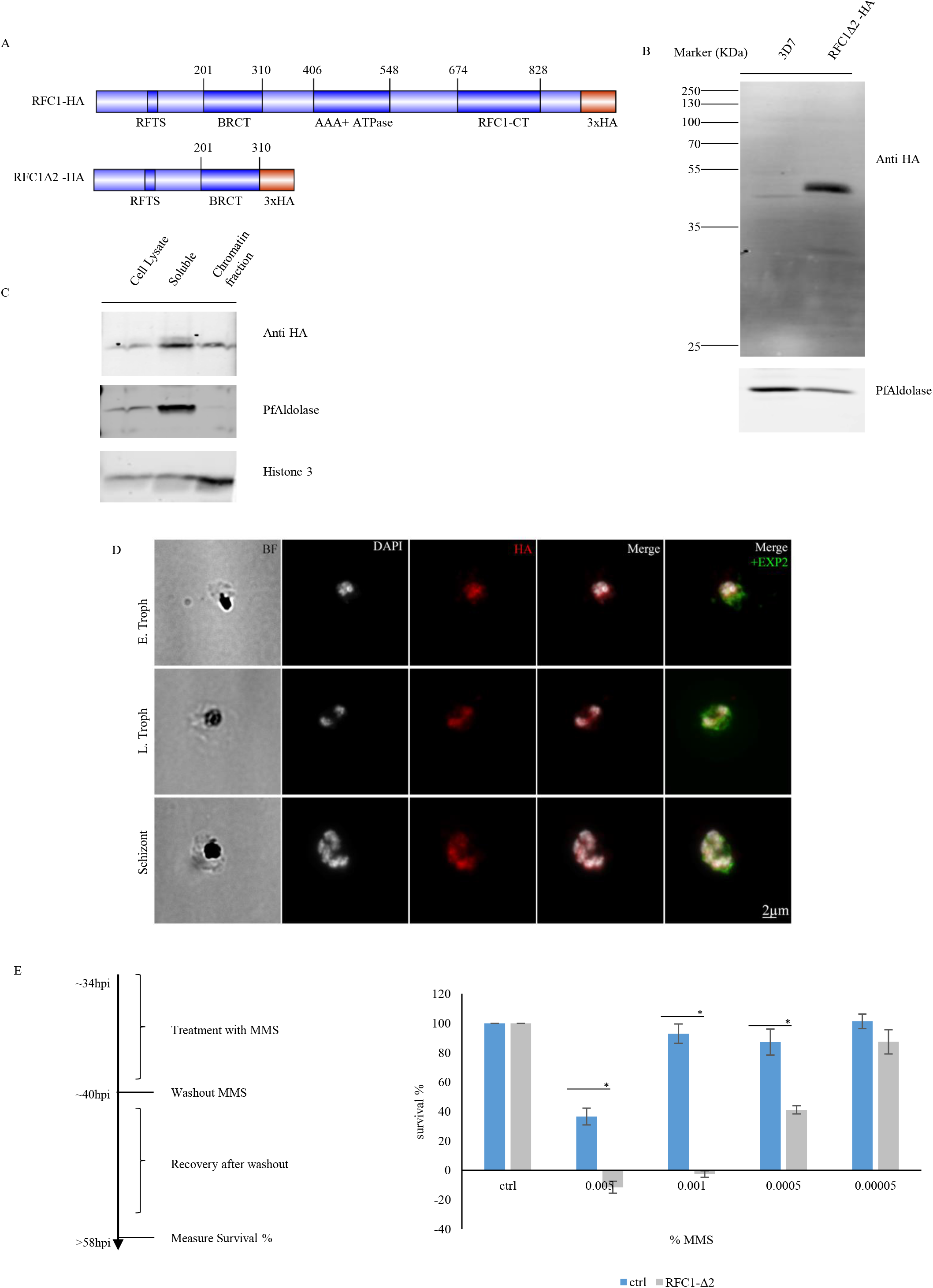
PfRFC1Δ2 -HA affects DNA damage recovery. (A) A cartoon representation of the domain positions in the full length RFC1 protein as well as that of the N-terminal truncation containing the RFTS and the BRCT domain. (B) The parasite lysates from the wild type 3D7 and the RFC1Δ2 -HA were subjected to western blotting and probed with anti-HA antibody and PfAldolase as a loading control. Ectopically expressed PfRFC1Δ2-HA was observed at the expected size using anti-HA antibody. (C) RFC1Δ2-HA cells were subjected to fractionation using detergents to separate the cells into a soluble fraction and an insoluble chromatin bound fraction. Anti-HA antibody was used to detect PfRFC1Δ2-HA, and PfAldolase and PfHistone3 were used as controls. The detergent resistant, PFAldolase free fraction identified the presence of PfRFC1Δ2-HA confirming its interaction with chromatin. (D) Immunofluorescence assay was performed on the intraerythrocytic stages of PfRFC1Δ2-HA and probed with anti HA (Red), Anti-EXP2 (Green) and stained the nucleus with DAPI (White). PfRFC1Δ2-HA was present within the nucleus as diffuse signals in early and late trophozoites, and schizonts. (E) An assay to measure recovery from Genotoxic stress was performed as described in the flowchart. Trophozoite stage of PfRFC1Δ2-HA (~34Hpi) were subjected to DMSO as control or MMS treatments at various concentrations for 6 hours. The drug was then washed out and the parasites were allowed to recover and measurements were made in the next cycle. It was observed that PfRFC1Δ2-HA subjected to intermediate concentrations of MMS recovered to lesser extent (n=3, * represents P<0.05) than the vector control parasites.

It has been suggested that the DNA repair continues upon washout of genotoxic stressors. Our data suggests that the truncated protein localizes in the nucleus and associates with DNA potentially providing us with a tool to evaluate the ability of the parasite to deal with DNA damage upon overexpression of a fragment incapable of forming the RFC complex. PfRFC1Δ2-HA contains the RFTS known to be involved in PCNA binding, a protein critical in the DNA repair pathway. The ability of PfRFC1 Δ2-HA to interfere with the DNA repair in the presence of the full length native PfRFC1 was therefore investigated (Fig. 4E). Recovery upon washout of the genotoxic drug MMS from PfRFC1Δ2-HA expressing parasites treated with 0.005%, 0.001%,and 0.0005%of MMS showed significant lower survival rates than the control, suggesting hampered DNA damage repair. A parasite line transfected with the plasmid containing only the HA tag under the same drug pressure was utilized as control and these recovered similar to PfRFC1Δ2-HA parasites under DMSO treatment and treatment with 0.00005% MMS. These results suggest that the presence of the N-terminal domain containing both the PCNA binding motif as well as the BRCT domain of PfRFC1 interferes with the DNA repair mechanisms likely due to the formation of non-functional DNA repair complexes.

### Effect of antimalarials on PfRFC1

As numerous antimalarial drugs such as artemisinin and chloroquine also induce genotoxic stress leading to cell death^1,46^, we were interested in establishing the role of PfRFC1 in responding to these types of stressors. The trophozoite stages of PfRFC1-3xHA were treated for 6 hours independently with various concentrations of Artesunate (ART) as well as chloroquine (CQ). sodium azide was used as an agent to induce necrosis. ART treatment of the parasites was performed at concentrations of 1x IC_50_ (2 nM) and 10x IC_50_ (20nM) which were reported to induce DNA fragmentation in trophozoite stages at 1 hour post treatment^2^. CQ treatment was performed at 1x IC_50_ (30 nM), 10x IC_50_ (300 nM), 100x IC_50_ (3 μM) concentrations. At low concentrations of CQ (1x, 10x IC_50_) DNA damage was previously reported with markers of apoptosis being activated only at higher concentrations (100x IC_50_)^1^. Western blotting of parasite lysates obtained after the treatments showed no significant change for 2 nM ART treatment while a small reduction of PfRFC1 levels at 20 nM of ART treatment was observed (Fig. 5A). Similarly, the highest concentration of CQ led to a significant reduction of the protein (Fig. 5A). The parasites subjected to high concentrations of sodium azide (1%) treatment showed a significant reduction of full length PfRFC1 arising from unregulated necrosis (Fig. 5A). The significant reduction of PfRFC1 observed (Fig. 5A) is a deviation from the up-regulation of PfRFC1 observed upon MMS treatment (Fig. 3A,B) and is possibly the result of protease activity induced at high drug concentrations leading to apoptosis / programmed cell death like effects on PfRFC1. This is in agreement with the observation that high levels of CQ leads to activation of apoptosis like features observed via the cleavage of DEVD/G motifs by proteases^1^. PfRFC1 contains a conserved DEVD/G motif in box V and this motif has been found to be cleaved upon induction of apoptosis in a variety of cell types^20^.

**Fig. 5:**
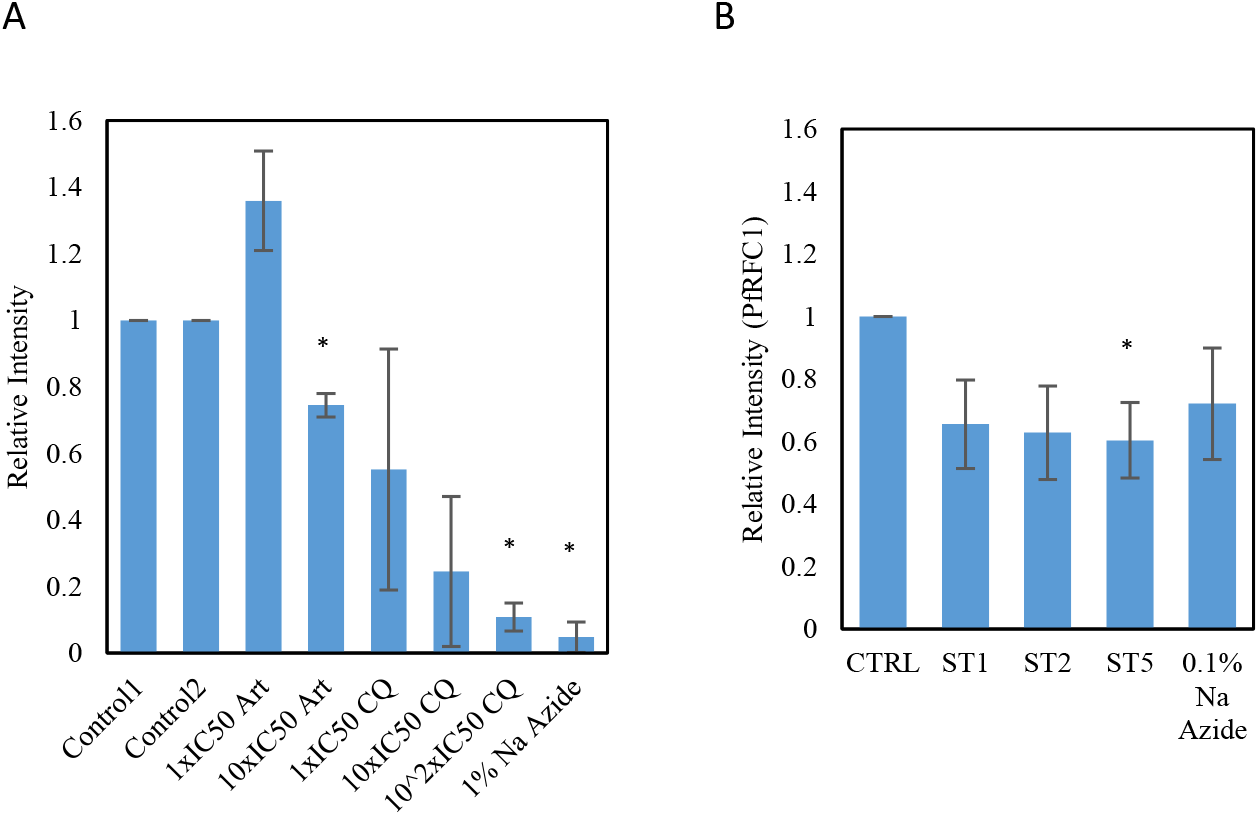
Effect of antimalarial drugs on PfRFC1. **(A)** A bar chart representing the levels of PfRFC1 upon treatment of PfRFC1-3HA trophozoite parasites with controls, Artesunate at 1xIC_50_, 10xIC_50_, and Chloroquine at 1xIC_50_, 10xIC_50_ and 100xIC_50_ for 6 hours. 1% Sodium azide treated parasites were used as a control for unregulated necrosis. Parasites treated with 10xIC_50_ Artesunate and 100xIC_50_ of Chloroquine showed significant reduction in the levels of full length PfRFC1 comparable to that of 1 % Sodium azide treated parasites. (n=3, * represents p<0.05). **(B)** A bar chart representing the levels of PfRFC1 upon treatment of synchronized trophozoite parasites expressing PfRFC1-3HA with staurosporine (ST) at 1, 2, 5 μM. (n=3, * represents p<0.05). The levels of PfRFC1 reduced significantly at 5 μM ST. 0.1% sodium azide, a necrosis control showed no significant change.

We were therefore interested in identifying the contribution of apoptosis/programmed cell death (PCD) like features on PfRFC1 upon treatment with antimalarial drugs. In order to measure the effect of apoptosis on PfRFC1 protein levels, concentrations of Staurosporine (ST) at 1, 2, 5 μM were added to synchronized trophozoites. These parasites were treated for 10 hours and 0.1% sodium azide was used as a necrosis control. The signals of PfRFC1 significantly reduced to 60.32% as compared to control amounts at Staurosporine (ST) concentration of 5 μM (Fig. 5B). This is lower than 10 μM ST used previously to induce apoptosis in *P.falciparum* under similar conditions^1^. No significant reduction was observed upon treatment with a necrosis control agent, 0.1% sodium azide. This indicated an apoptosis specific reduction of PfRFC1 levels. The parasites treated with 1% sodium azide as well as high doses of antimalarials therefore had a more significant apoptotic response towards PfRFC1 (Fig. 5A) leading to a reduction in the protein levels. The western blots with anti-HA antibody and PfAldolase as loading controls are provided for the various antimalarial and Staurosporine treatments (Fig. S3 A, B). PfRFC1Δ2-HA parasites overexpressing a truncated PfRFC1 lacking the DEVD/G protease cleavage site did not show a ST or necrosis dependent change in the protein levels as observe via western blotting (Fig. S3C).

### Mislocalisation of RFC1 is detrimental

As RFC1 was successfully tagged by modifying its endogenous loci, we utilized the knock sideways (KS) approach of functional analysis^29,47,48^. Here, PfRFC1 was endogenously tagged with 2xFKBP-GFP and also transfected with a plasma membrane mislocalizer containing FRB*-mCherry. The cell line, PfRFC1-GFP was validated for integration at the correct locus (Fig. 6A).The expression was verified by probing the cell lysate for GFP (Fig. 6B). Upon addition of 200 nM rapamycin, FRB* rapidly dimerizes with FKBP thereby sequestering PfRFC1 from the nucleus into the parasite plasma membrane. This was evident as early as 12 hours post addition of rapamycin where PfRFC1-GFP was observed to be present in the parasite cytoplasm (Fig. 6C).

**Fig. 6:**
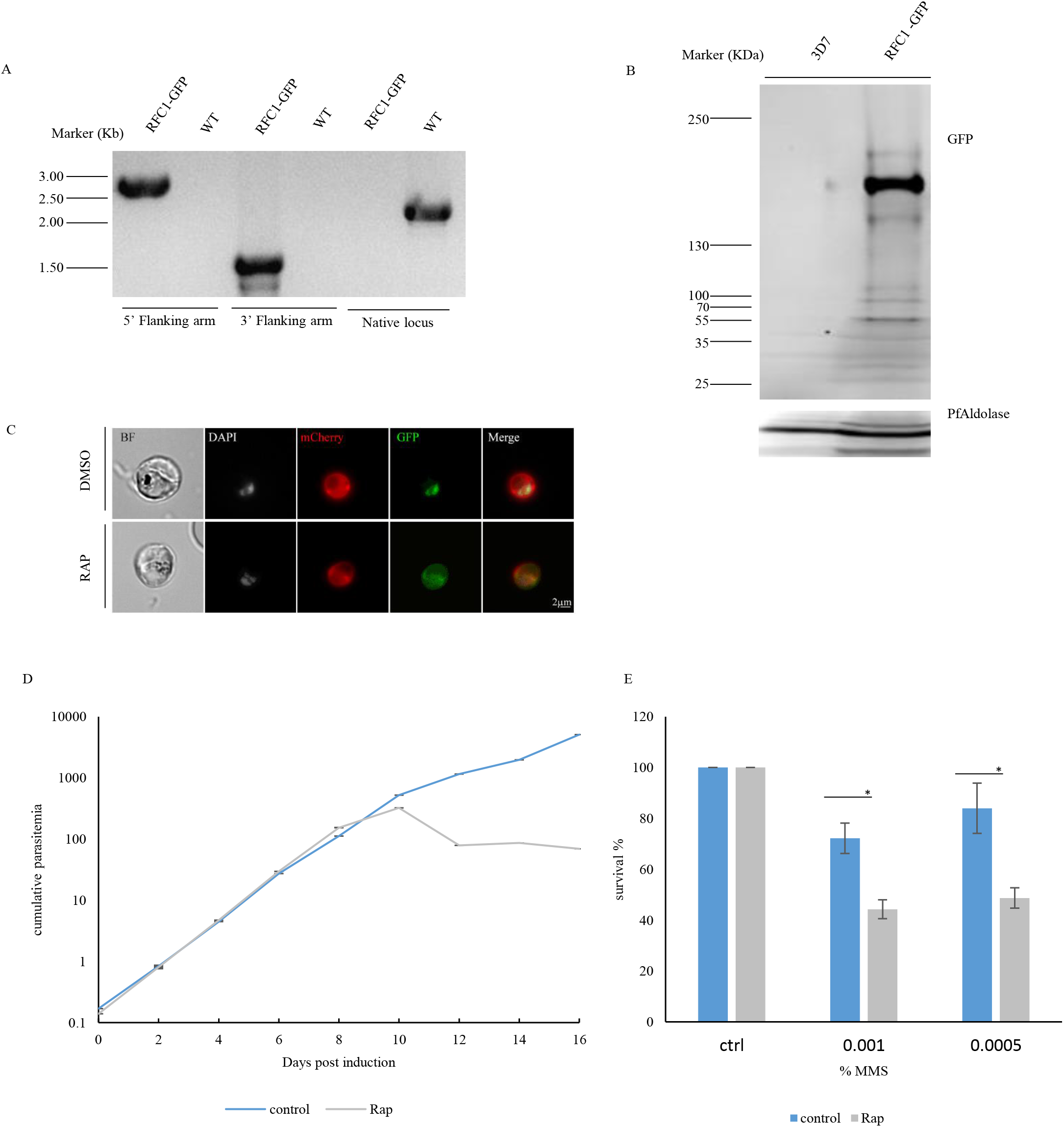
Mislocalisation of PfRFC1-GFP is detrimental. (A) PfRFC1 was endogenously tagged with a 2xFKBP-GFP tag and integration was verified via PCR. (B) The parasite lysates from the wild type 3D7 and the PfRFC1-GFP were subjected to western blotting and probed with anti-GFP antibody and PfAldolase as a loading control. The endogenously expressed PfRFC1-GFP was observed at the expected size using anti-GFP antibody. (C) Parasites treated with control or Rapamycin were observed under the fluorescence microscope. In control treated parasites PfRFC1-GFP (Green) was observed in the nuclear fraction while the mislocalizer was observed in the parasite periphery (Red). Upon treatment with Rapamycin, the GFP signals of PfRFC1-GFP were sequestered beyond the nucleus into the parasite. (D) The parasitemia was traced for PfRFC1-GFP parasites treated with control or rapamycin. On cycle 5 onwards the parasitemia in the treated parasites depreciated leading to death (n=3, * represents p<0.05). (E) A recovery from Genotoxic stress assay was performed on day 6 trophozoite stage of PfRFC1-GFP (~34Hpi) subjected to control or rapamycin induced mislocalization. These were treated with DMSO as control or MMS treatments at various concentrations for 6 hours. The drug was then washed out and the parasites were allowed to recover and measurements were made in the next cycle. It was observed that PfRFC1-GFP subjected to MMS stress recovered to lesser extent (marked with *,P<0.05) than the control treated and non-mislocalized parasites.

The parasites expressing the tagged PfRFC1 and the plasma membrane mislocalizer were treated with the inducer Rapamycin and a control. The parasitemia was tracked over a period of weeks (Fig. 6D). On Day 10 onwards, the cell line subjected to rapamycin treatment showed retardation in growth eventually leading to cell death highlighting the essentiality of PfRFC1. The ability of the cells to recover from DNA damage upon the specific mislocalization of PfRFC1 was investigated at day 6 where the control and mislocalized parasites continue to grow at comparable rates. The ability of these parasites to perform DNA repair upon MMS mediated genotoxic stress was also evaluated. Synchronized control and rapamycin treated trophozoites at day 6 were treated with 0.001% and 0.0005%, of MMS. DMSO was used as a control treatment. The parasites subjected to the mislocalizer rapamycin and treated with MMS recovered to a significantly lesser extent than the MMS treated and non-mislocalized parasites (Fig. 6E). This suggested that the ability to recover from DNA damage is perturbed in *Plasmodium* upon PfRFC1 mislocalization.

## Discussion

The involvement of DNA damage repair response in *P. falciparum* parasites treated with antimalarial drugs such as artemisinin has highlighted the need to understand this mechanism^22^. The emerging resistance to artemisinin involving delayed asexual growth stages indicates an involvement of the replication machinery^6^. Further, studies aimed at identifying the molecular response in parasites exposed to genotoxic stress have identified a variety of early transcribed genes^22^. PfRFC1 is an important component of the DNA repair pathway in these parasites and was observed to be upregulated upon genotoxic stress^22^.

The localization of the largest subunit of this complex PfRFC1 has been observed to be developmentally regulated. In the trophozoite stage where active DNA replication occurs, PfRFC1 is observed in punctate patterns within the nucleus. These foci are the probable regions of interactions with PfPCNA1 via the PCNA binding motif present in both the N and C terminal of PfRFC1.These replication foci contain the replication machinery and PCNA is considered a marker for these replication factory sites^24^. As the parasites mature into well segmented schizonts towards the end of replication, PfRFC1 is sequestered at the nuclear periphery (Fig. 1H). This is in agreement to studies on the spatial distribution of replication sites showing replication factories within the nucleus in S phase which translocate to the nuclear periphery towards the end of replication at sites where the telomeres are attached to the nuclear membrane^49^. The staining pattern of PfRFC1 with PfH3K4Me3 in late stage schizonts show specific regions of overlap at the nuclear periphery hinting at the presence of intergenic regions undergoing replication at the nuclear periphery as well (Fig. 3C). These PfRFC1 signals further translocate from the external nuclear periphery in segmented schizonts to a single punctae in the egressing merozoites (Fig. 1H). The localization of PfRFC1 therefore reflects a well-orchestrated progression of distinct patterns of replication in the developing merozoites. The change in localization post replication has been noted in the case of PfORC1, PfPCNA1 as well as PfORC5, however a clear external nuclear peripheral localization has not been documented^50,51^. PfORC1, the origin recognition complex protein 1 observed at the replication foci during S phase tends to be degraded at the segmented schizont stages^50^. Other well documented plasmodium replication proteins such as PfPCNA1 and PfORC5 are known to disassemble from the replication foci post replication^50^.

This PfRFC1 complex was effectively immunoprecipitated and confirmed via mass spectrometry and western blotting. Proteins enriched in the IP indicate the successful enrichment of the PfRFC1-5 complex (Fig. 2 A, B). The proteins identified are significantly enriched in oxidative response, DNA repair and damage recovery proteins. PfAlba1 was notably enriched. PfAlba1 has been previously described to be a perinuclear protein in the ring stages, migrating to the cytoplasm in the mature intra erythrocytic stages of the parasite^52^. The Significance of the PfRFC1 complex and any potential interactions with PfAlba1 remains to be determined.

The survival of the plasmodium parasite within the host depends on its ability to counteract hostile environmental threats such as the host immune response, oxidative free radicals and drug challenges. The parasite has numerous mechanisms of protecting its DNA such as the Base excision repair, nucleotide excision repair, homology independent end joining mechanisms, and the Rad51 mediated homologous recombination pathway^22–24^. This study elaborates on the RFC1 protein, a factor involved in DNA replication and nucleotide excision repair^53^.When the replicating trophozoite stage parasites were subjected to genotoxic stress via differentially acting drugs such as MMS and HU, a robust accumulation of PfRFC1 was observed in the cell lysates (Fig. 3 A, B). The increase in protein levels is also reflected in the increase in levels of the soluble and chromatin bound PfRFC1 in the parasite nucleus (Fig. S2 A, B). Additionally, the Perinuclear localization of PfRFC1 observed in segmented schizonts stages reflecting a distinct pattern of progression of replication is affected upon genotoxic stress leading to the enrichment of PfRFC1 into numerous distinct punctae within the nucleus (Fig. 3C). The sites containing PfRFC1 retained within the nucleus potentially harbor other repair components such as PfPCNA1, PfORC5 etc. in cells subjected to genotoxic stress. The functional increase in the level of the chromatin bound form has been associated with DNA repair activity mediated by polδ or Polε^54^ Numerous excision repair components such as FEN-1, DNA Ligase 1 etc. have also been identified in *P.falciparum^55^* and the involvement of PfPCNA1, 2 and PfFEN-1 has been studied to be key in the long patch BER^56^. This increase in replication machinery and recombination repair components upon DNA damage has been observed in *P.falciparum* for proteins such as PfRFC1, PfPCNA1, PfRAD51 and PfFEN-1^22,24^.

PfRFC1 digresses from reported RFC1 proteins via the elongated N-terminus segment with a recessed replication factory targeting sequence (Fig. 1B, C). This region isn’t essential for viability in human and yet it plays an important role in vivo to facilitate DNA damage repair^18^. An N-terminal region containing the RFTS, a PCNA binding domain, and the BRCT domain was found to be generated during apoptosis and localizes to sites of DNA damage by interacting with PCNA^57^. Additionally, in a variety of cell types, RFC1 has been reported to be cleaved by caspase-3 at an evolutionarily conserved motif (DEVD/G) spanning Box V-VI upon activation of apoptosis^20,21^. This Caspase-3 generated N-terminal fragment actively inhibits DNA replication, thereby mediating cell cycle arrest. An assay designed to measure recovery of parasites from genotoxic stress indicated that the ectopic expression of a similar N-Terminal fragment of PfRFC1 leads to reduced parasite survival only upon DNA damage (Fig. 4D). *P. falciparum* lacks molecular evidences for pathways leading to apoptosis or programmed cell death (PCD). The absence of caspase homologs in *P. falciparum* hints at the involvement of metacaspace orthologs or clan CA/CD cysteine proteases. Studies using Chloroquine (CQ) treated parasites have recorded the induction of DNA fragmentation, activation of cysteine proteases, and features of PCD^1,58^. Artesunate (ART) a derivative of artemisinin induced DNA double strand breaks in *P. falciparum* leading to the generation of reactive oxygen species ultimately resulting in cell death^2^. Therefore, the effect of Antimalarial drugs such as artemisinin and chloroquine (CQ) on PfRFC1 was investigated. The parasites subjected to high concentrations of CQ and ART, resulted in reduced levels of full length PfRFC1 unlike the control or low dose treatments (Fig. 5 A, B). Additionally, comparing the treatment of various concentrations of an apoptotic agent staurosporine highlighted an apoptosis specific reduction in the levels of full length PfRFC1 at 5μM ST lower than the 10 μM previously used to observe apoptosis in *Plasmodium^1^*.

The mislocalization of the endogenous PfRFC1 from the nucleus is associated with cell death at day 10 post treatment (Fig. 6 C, D). The delayed effect on cell growth could be attributed to the levels of the mislocalizer and the number of FKBP domains on the RFC1 protein as observed with other parasite nuclear proteins subjected to knock-sideways^29^. Additionally, the progressive mislocalization of PfRFC1 over numerous cell cycles could have compounded its effect leading to a growth inhibition phenotype. The cells subjected to mislocalization also presented a defect in recovery from genotoxic stress earlier than the measurable growth effect (Fig. 6 E). These results suggest an essential role for PfRFC1 in responding to genotoxic stress in addition to its replication function.

We propose a model (Fig. 7) where an orchestrated sequence of replication events occur in trophozoite stages at the nucleus marked by punctate regions leading to DNA synthesis. These replication foci contain other replisome components such as PfPCNA1, PfORCs etc. and completion of replication in the schizont stages leads to their disassembly.PfRFC1 migrates to the nuclear periphery in schizonts and further outside the nucleus as a punctae in merozoites. Genotoxic stress however, affects this natural progression of the replication cycle leading to the recruitment of repair components to sites of DNA damage as evidenced by the absence of PfRFC1 at the nuclear periphery upon genotoxic stress. This study highlights the interplay between replication progression and DNA damage and recovery signals contributing to cell death.

**Fig. 7:**
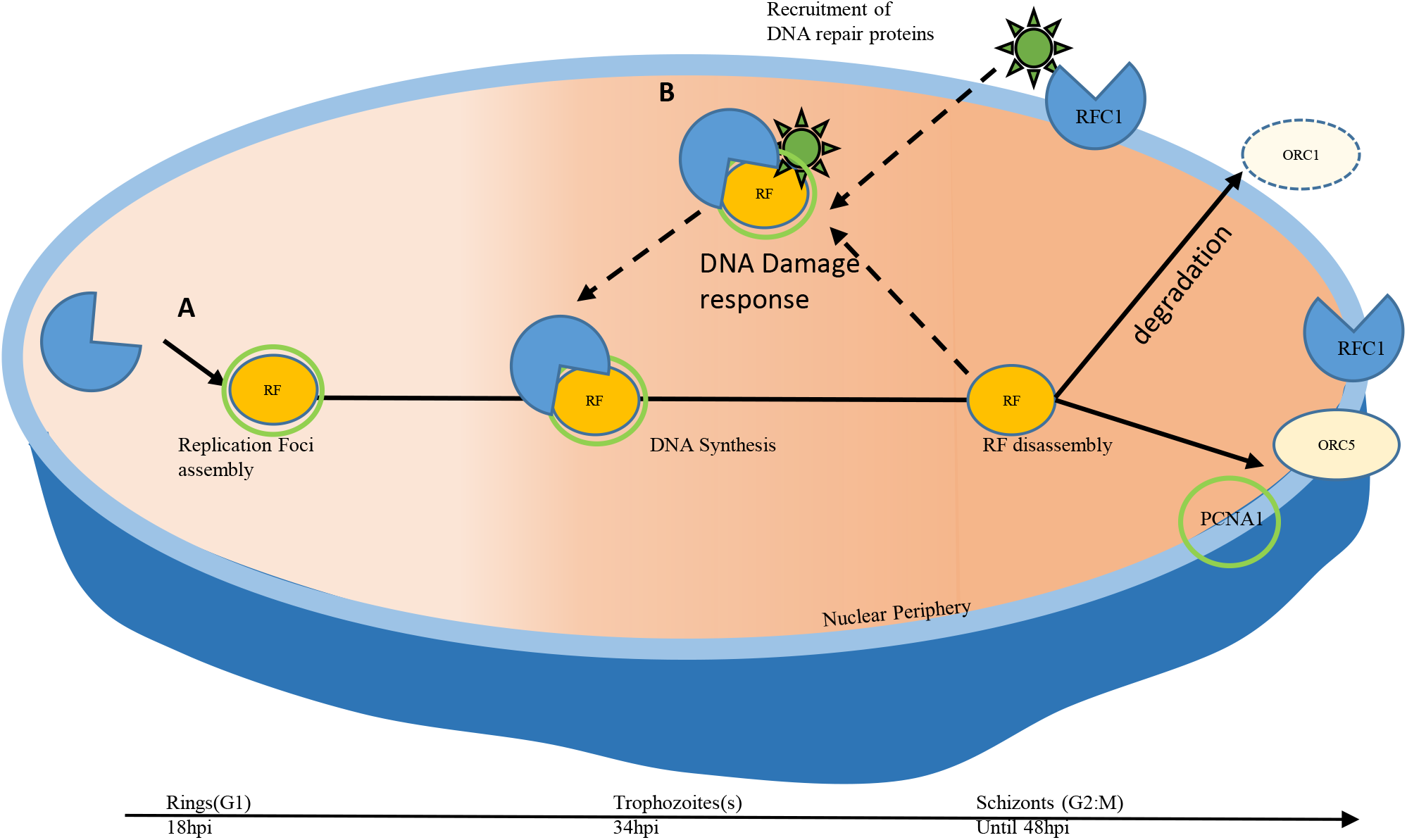
model representing the role of PfRFC1 in replication and DNA damage repair of *P. falciparum:* **(A)** The replication foci begins to form in the maturing ring stages of the parasites and PfRFC1 is recruited to these sites via potential interactions with other replisome proteins such as PfPCNA1. The Replication foci helps synthesize new DNA thereby generating new DNA content for the maturing trophozoites and schizonts. Upon completion of replication, while PfPCNA1 and PfORC5 are disassembled from the replication foci, PfRFC1 is sequestered to the nuclear periphery. (**B**) When the Parasites are subjected to genomic stress PfRFC1 and other DNA repair proteins are recruited to the sites of DNA damage promoting its repair. This leads to the presence of PfRFC1 within the nucleus and potential halting of DNA replication.

## Acknowledgement

WR99210 used was a kind donation from Jacobus Pharmaceuticals, Princeton, NJ. pSLI-2xFKBP-GFP and pLyn-FRB-mCherry-nmd3-BSD were a gift from Tobias Spielmann. We thank Prof. Suman Kumar Dhar, Special Centre for Molecular Medicine of Jawaharlal Nehru University, India for his kind assistance with the PCNA1 antibody western blot. The authors are grateful to the members of Peter Preiser’s lab for the critical reading of the manuscript. This research is supported by the Singapore Ministry of Health’s National Medical Research Council under its Open Fund Individual Research Grants (OFIRG17may073), the Singapore Ministry of Health’s National Medical Research Council under its Cooperative Basic Research Grant (CBRG12nov014) and the Singapore Ministry of Education, Singapore, under the NTUitive Gap Fund (NGF-2017-03-032). The funders had no role in study design, data collection and interpretation, or the decision to submit the work for publication.

## Supplementary Data

**Fig. S1:**
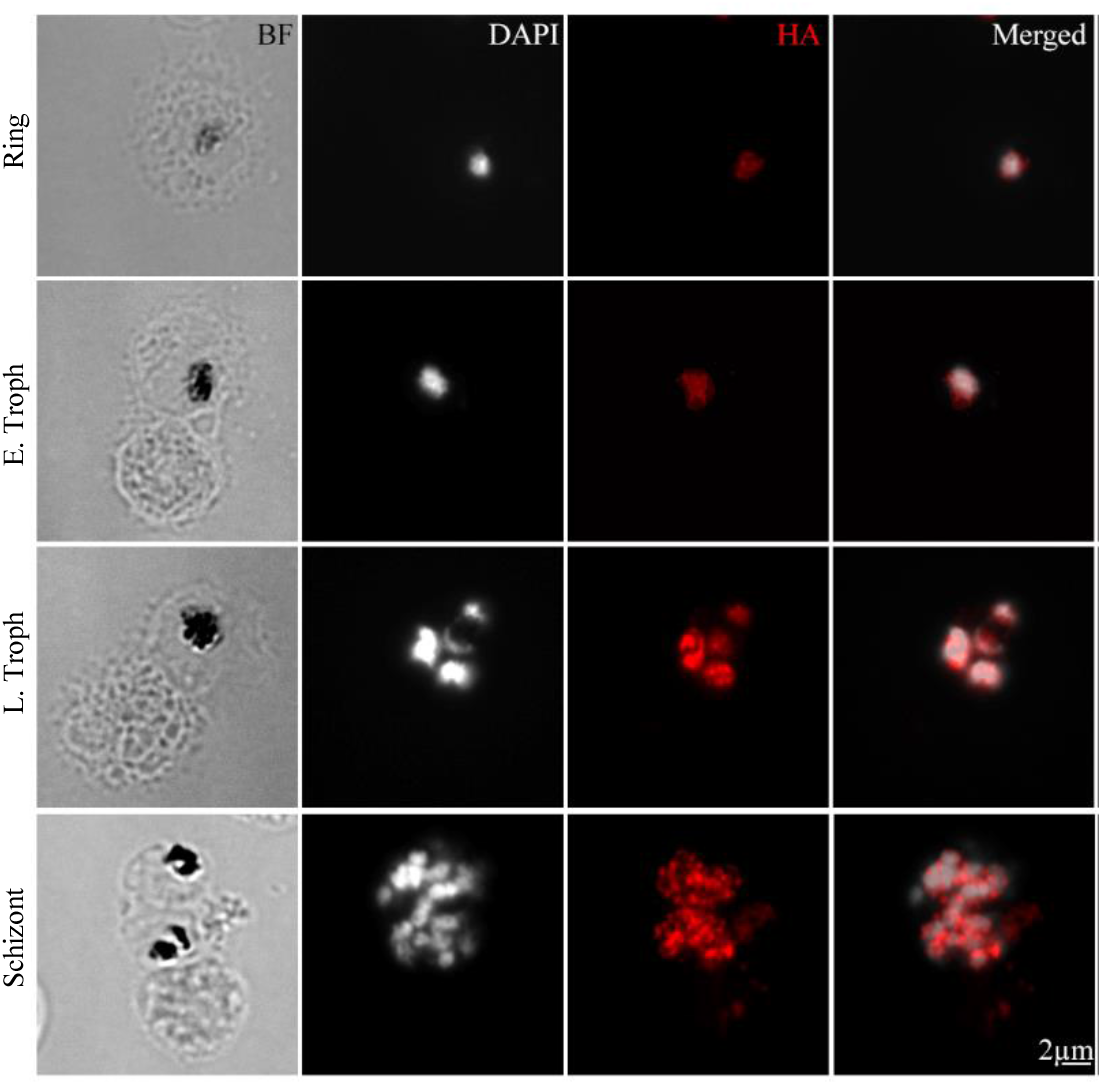
PfRFC1 localizes dynamically throughout intraerythrocytic developmental cycle: IFA was performed on smears of tagged PfRFC1 expressing parasites and probed with anti-HA antibody (Red) to mark PfRFC1 which localizes at the nucleus stained by DAPI (White). The ring, early and late trophozoite, and schizonts were identified using bright field (BF) and by the nuclear staining.

**Fig. S2:**
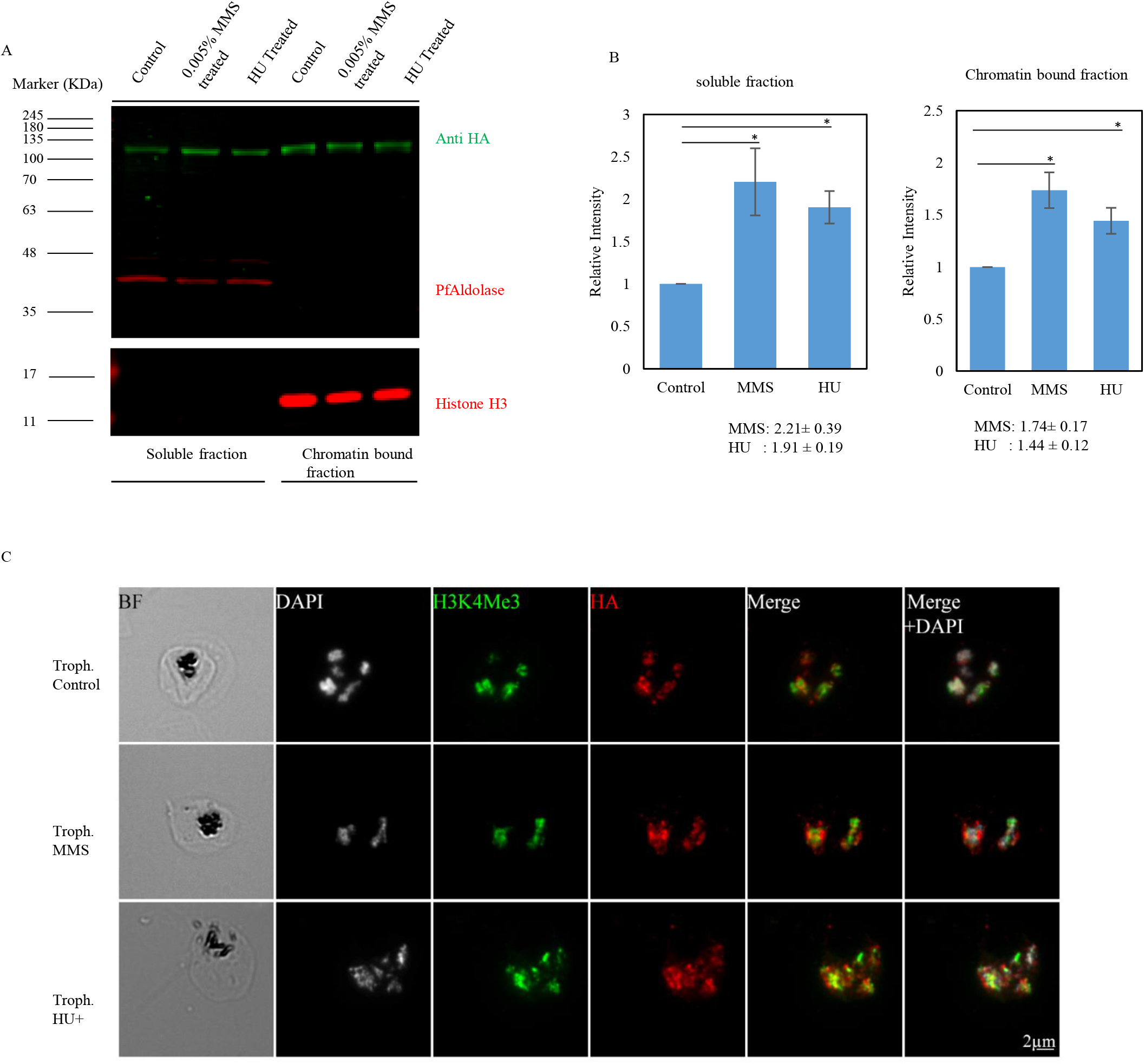
PfRFC1 is stimulated upon DNA damage. PfRFC1-3HA parasites treated with MMS and HU were extracted into a detergent soluble fraction and an insoluble (IN) or the chromatin bound fraction (SF) and analyzed by western blotting. PfRFC1 was detected using anti-HA antibody. PfAldolase was used as a loading control for the soluble fraction and PfHistone3 for the chromatin bound fraction. (B) Quantification of the various samples via the normalization of the PfRFC1 signals with regards to PfAldolase or PfHistone3 revealed a significant upregulation in the samples treated with MMS or HU. The results show means ±S.E.M (n=3, IN: p=0.013 for MMS and p=0.023 for HU; SF: p=0.038 for MMS and p=0.009 for HU). **(C)** Cells found at the trophozoite stages after the 6 hour treatment were also subjected to immunofluorescence and probed with anti HA (Red), Anti H3K4Me3 (Green) and stained the nucleus with DAPI (White). The nuclear localization of PfRFC1 remained comparable irrespective of the treatments at the trophozoite stages.

**Fig. S3:**
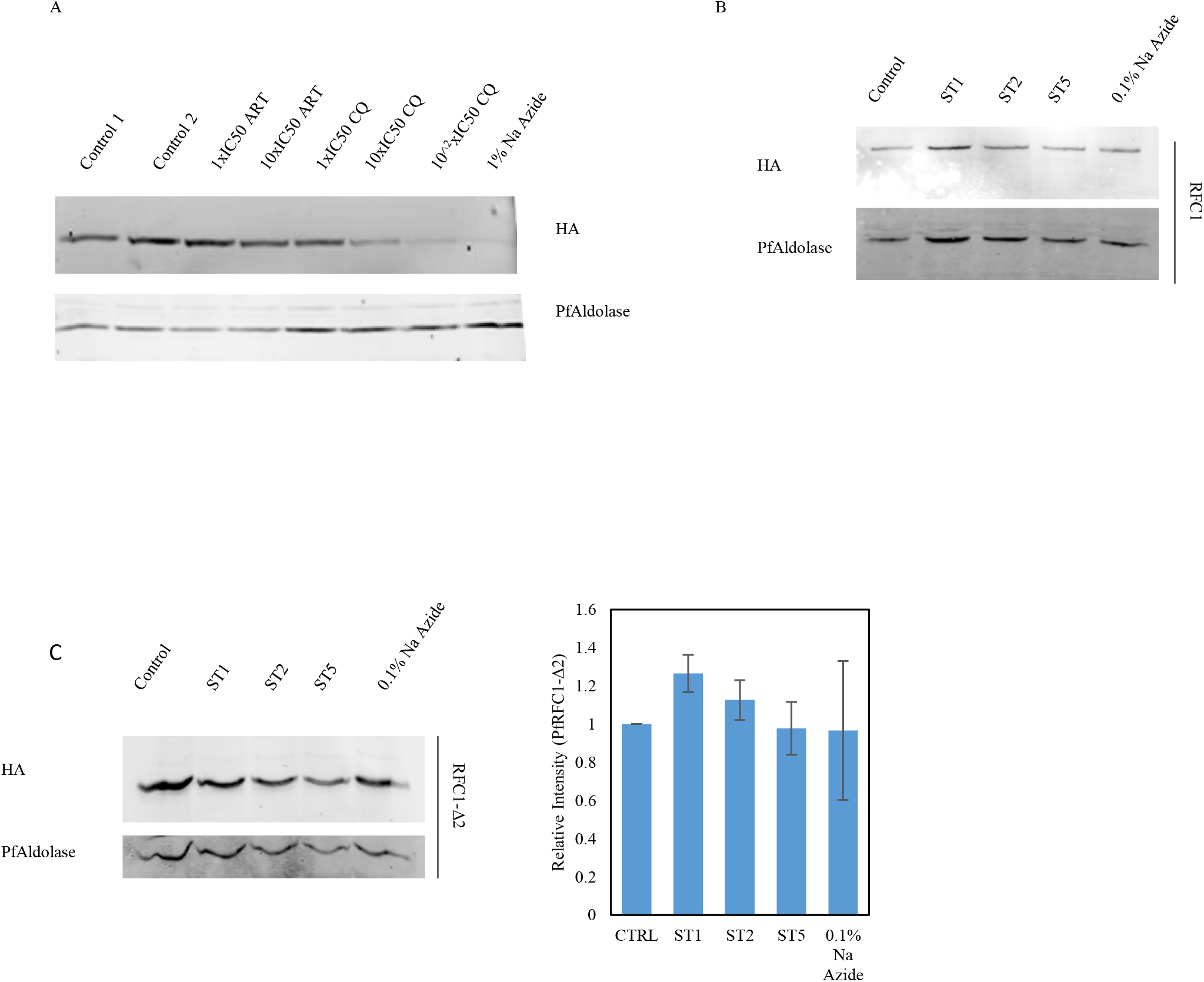
Effect of antimalarial drugs on PfRFC1: **(A)** PfRFC1-3HA trophozoite parasites were treated with controls, Artesunate at 1xIC_50_, 10xIC_50_ and Chloroquine at 1xIC_50_ 10xIC_50_ and 100xIC_50_ for 6 hours. 1% sodium azide treated parasites were used as a control for necrosis. Parasites treated with 10xIC_50_ Artesunate and 100xIC_50_ of Chloroquine showed significant reduction in the levels of full length PfRFC1 comparable to that of 1% Sodium azide treated parasites. Anti-HA antibody detected PfRFC1. PfAldolase was used as a control protein and its levels remains unchanged. **(B)** Synchronized trophozoite PfRFC1-HA parasites were treated with staurosporine (ST) at 1, 2, 5 μM. The levels of PfRFC1 reduced significantly at 5 μM ST. 0.1% sodium azide, a necrosis control showed no significant change. Anti-HA antibody detected PfRFC1. PfAldolase was used as a control protein and its levels remains unchanged. **(C)** Staurosporine (ST) at 1, 2, 5 μM were added to synchronized trophozoite parasites expressing PfRFC1Δ2-HA. These parasites were treated for 10 hours and 0.1% sodium azide was used as a necrosis control. The parasites were released via saponin treatment and subjected to western blotting using anti-HA antibody to detect PfRFC1Δ2. PfAldolase was used as a control protein and its levels remains unchanged. The bar chart summarizes the levels of PfRFC1Δ2 upon treatment with staurosporine (ST) at 1, 2, 5 μM. (n=3). The levels of PfRFC1Δ2-HA did not alter significantly. 0.1% sodium azide, a necrosis control showed no significant change.

**Table S1:** List of all primers used in the study.

**Table S2:** List of all P.falciparum and contaminating proteins detected via mass spectrometry.

